# Learning the All-Atom Equilibrium Distribution of Biomolecular Interactions at Scale

**DOI:** 10.64898/2026.03.10.710952

**Authors:** Yusong Wang, Youjun Xu, Wentao Li, Haoyu Yu, Wenjuan Tan, Shaoning Li, Qiaojing Huang, Nanjun Chen, Xuan Wu, Qilong Wu, Kai Liu

## Abstract

Biomolecular functions are governed by dynamic conformational ensembles rather than static structures. While models like AlphaFold have revolutionized static structure prediction, accurately capturing the equilibrium distribution of all-atom biomolecular interactions remains a significant challenge due to the high computational cost of molecular dynamics (MD). We present AnewSampling, a transferable generative foundation framework designed for the high-fidelity sampling of all-atom equilibrium distributions, which is the first model to faithfully reproduce MD at the all-atom level. It uses a quotient-space generative framework to ensure mathematical consistency and leverages the largest self-curated database of protein-ligand trajectories to date, with over 15 million conformations. Statistically, AnewSampling consistently outperforms all prior generative methods on the ATLAS monomer benchmark, and the all-atom capabilities of AnewSampling enable close statistical alignment with ground-truth MD for evaluating atomic biomolecular interactions in protein-ligand dynamics. Furthermore, AnewSampling successfully recovers coupled ligand and side-chain motions in CDK2 systems, overcoming a major sampling hurdle inherent to conventional MD. AnewSampling enables rapid exploration of conformational landscapes prior to intensive simulations, elucidating fundamental biophysical mechanisms and accelerating the broader design of functional biomolecules.

## 1 Introduction

Biomolecular functions are fundamentally dictated by precise, atomic-scale physical interactions between molecules, which constitute the essential machinery of life and drive complex processes ranging from enzymatic catalysis [1] to intricate cellular signaling [2]. Consequently, achieving a comprehensive understanding of the biophysical mechanisms underlying these interactions, alongside developing robust methods for their artificial modulation, represents one of the most significant challenges in contemporary science. Notably, the structural characterization of these atomic interfaces has recently undergone a revolutionary transformation, largely driven by geometric deep learning architectures such as AlphaFold [3–6] and related predictive models [7–11]. While it is now possible to predict the static ground states of biomolecular assemblies with unprecedented accuracy, these rigid snapshots offer only a limited perspective. In physiological environments, biomolecules exist in a state of constant conformational fluctuation, where biological efficacy is frequently governed by the dynamic transitions between diverse structural states rather than the geometry of a single structure [12]. Therefore, to comprehensively understand and engineer molecular function, it is imperative to move beyond static structural descriptions and accurately recover the dynamic equilibrium dictated by the Boltzmann distribution.

Experimental techniques such as cryogenic electron microscopy (cryo-EM) can resolve diverse conformational states [13], yet their application remains limited by high costs and the intensive time required for sample preparation and data collection. Molecular dynamics (MD) simulations [14] provide a more accessible alternative for characterizing these fluctuations at atomic resolution. While MD has long been the standard for studying biomolecular dynamics, the serial nature of numerical integration requires femtosecond-scale timesteps, creating a major computational bottleneck. Even relatively simple protein folding phenomena, which typically occur on microsecond-to-millisecond scales, can currently only be captured using special-purpose supercomputers [15]. Since such specialized hardware is not widely accessible for routine drug discovery, the dependence on conventional physical simulations ultimately limits the throughput and scalability of pipelines that require a detailed understanding of molecular dynamics.

Recent advances in artificial intelligence (AI) have explored multiple avenues to overcome the inherent bottlenecks of molecular dynamics. One prominent strategy involves multiple sequence alignment (MSA) perturbation [16–18]. While these heuristic methods possess the capability to sample alternative conformations, they offer limited controllability and fail to reflect rigorous thermodynamic distributions. Concurrently, generative AI has begun to bridge this gap, demonstrating that deep learning can emulate MD simulations orders of magnitude faster. These emerging generative approaches generally follow two distinct paradigms: trajectory-based transition modeling and equilibrium distribution learning. The first paradigm focuses on learning transition operators to generate kinetic trajectories from an initial frame. Although these methods have demonstrated the capability for capturing free energy landscapes [19–21] and observe specific dynamical phenomena [22–24], they face significant challenges regarding sampling reliability. Due to the inherent autocorrelation between sequential samples, ensuring statistical convergence remains difficult. Consequently, the resulting ensemble may be sensitive to the choice of the initial frames. In parallel, another paradigm [25–27] seeks to directly learn transferable equilibrium distributions. While BioEmu [27] has emerged as a landmark framework in this category by adapting an architecture similar to AlphaFold2, it frequently relies on simplified chemical representations. By utilizing backbone-based SE(3) diffusion, it models only the distributions of the backbone while ignoring side chains and ligands, thereby stripping away the atomic interactions that drive drug binding. In the post-AlphaFold2 era, structure prediction [6] has shifted toward generative all-atom modeling; however, the scarcity of MD training data often limits the resulting structural diversity. While MD-enhanced models, such as Boltz2 [10], produce more distinct conformations, the mathematical inconsistencies between training objectives and sampling procedures in their generative frameworks hinder their ability to accurately recover true equilibrium ensembles. Moreover, the practical utility of all-atom ensemble emulators remains difficult to quantify, given the lack of consensus on standard evaluation metrics for characterizing the dynamics of biomolecular interactions [28]. These limitations highlight a fundamental question: *Can generative frameworks investigate biomolecular interactions at an all-atom level while faithfully reproducing converged MD equilibrium distributions in a transferable manner, and what metrics can provide a rigorous evaluation of these atomic interactions?*

Toward this end, we introduce AnewSampling, a foundational generative framework developed to sample from the all-atom equilibrium distributions of biomolecular complexes. As shown in Fig. 1b, by adopting a generative framework [29] that mathematically ensures consistency between training and sampling, we employ a sophisticated hybrid fine-tuning strategy to shift the objective from static structure prediction to stochastic dynamic modeling. The transferability of AnewSampling is underpinned by the construction of AnewSampling-DB (Fig. 1c), the largest database of protein-ligand dynamics to date, which encompasses over 15 million conformations. Spanning 10,297 unique protein sequences and 27,979 unique ligand SMILES, AnewSampling-DB covers a vast chemical space of both congeneric and non-congeneric series. All trajectories are generated through a unified pipeline using a consistent set of force fields to ensure rigorous physical consistency across diverse interaction profiles. Furthermore, we address the fundamental ambiguity of evaluating generative dynamics by proposing a multi-level assessment strategy (Fig. 1a). This framework rigorously validates physical fidelity by quantifying ligand conformations, protein-ligand interaction patterns, and intrinsic protein dynamics.

**Figure 1.**
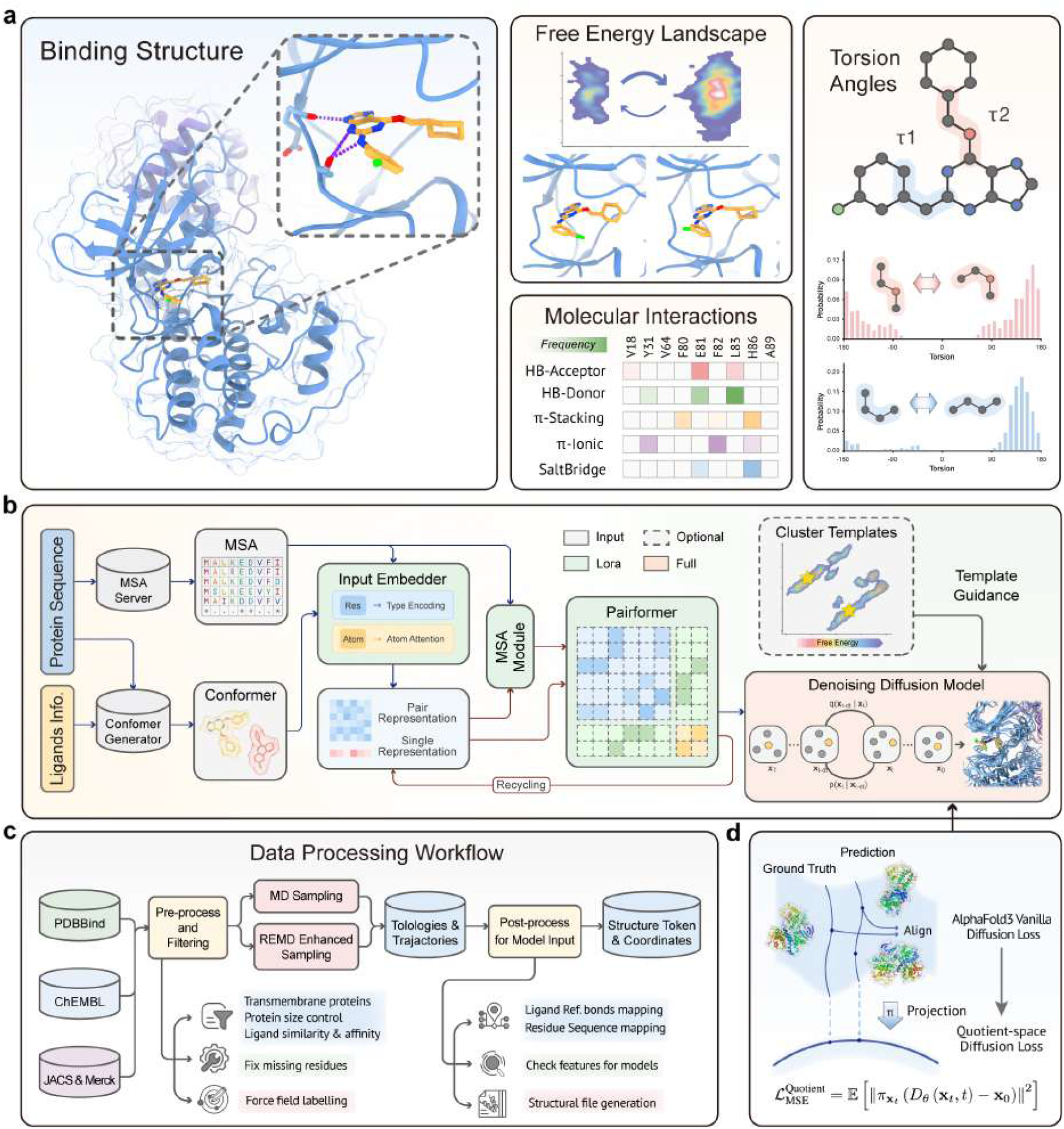
Overview of the AnewSampling framework for sampling biomolecular conformational ensembles. **(a)** Characterization of the dynamic binding landscape. Beyond a single static structure, protein-ligand interactions are visualized via free energy landscapes with distinct metastable states. These are further characterized by dynamic molecular interaction fingerprints and multimodal torsion angle distributions. **(b)** AnewSampling architecture and training objective. The model processes protein sequences (via MSA) and ligands through the Input Embedder, MSA Module, and Pairformer module. To shift the objective from static structure prediction to stochastic dynamic modeling, Low-Rank Adaptation (LoRA) is integrated into the sequence feature representation modules (green), while the Diffusion module undergoes full-parameter fine-tuning (red). The generation process is guided by clustered noisy templates, with training supervised by a quotient-space diffusion loss [29] 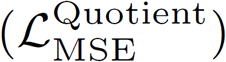 that projects coordinates onto a quotient manifold to resolve symmetry invariances. **(c)** Data curation and processing workflow. Raw structural data from PDBBind, ChEMBL, and JACS & Merck undergo rigorous quality filtering—including resolution truncation and redundancy removal—prior to MD and REMD enhanced sampling. The resulting trajectories are featurized and tokenized to serve as high-fidelity inputs for model training and testing.

We rigorously validated AnewSampling, beginning with the ATLAS [30] dataset of protein monomers to establish baseline performance. While existing generative methods are restricted to monomeric systems, AnewSampling consistently outperforms these specialized models across 13/13 evaluation metrics, confirming the fundamental efficacy and robustness of our generative framework. To assess protein-ligand dynamics, we utilized three distinct test sets: a diverse held-out set of PDB systems to demonstrate broad chemical generalizability, the JACS & Merck [31] datasets—industrial standards for structure-activity relationship (SAR) analysis—to capture subtle ligand-induced conformational changes and an in-house dataset derived from our internal drug discovery pipelines. Across these challenging tasks, AnewSampling achieves a statistical alignment with ground-truth MD distributions that far surpasses both pure static predictors and recent MD-enhanced models such as Boltz2 [10], which frequently suffer from mode collapse or fail to recover the full breadth of the equilibrium ensemble. Notably, while AnewSampling is trained exclusively on standard molecular dynamics trajectories, it demonstrates an emergent capability for enhanced sampling. By learning the underlying biophysical interactions shared across diverse chemical systems, the model effectively bypasses the high energy barriers that typically trap conventional MD. For CDK2 complexes (1H1R and 1H1S) [32], AnewSampling successfully navigates complex free energy landscapes to reproduce the multimodal ensembles observed in replica-exchange MD (REMD), recovering coupled ligand and side-chain motions that are often missed by conventional MD within practical simulation budgets.

The unprecedented computational efficiency of AnewSampling enables the rapid exploring of complex conformational landscapes, thereby facilitating the integration of the framework into diverse research and industrial pipelines. By providing a scalable method for the exploration of dynamic distributions prior to intensive physical simulations, AnewSampling catalyzes a paradigm shift toward dynamics-aware design, wherein the comprehensive characterization of biomolecular flexibility directly informs the development of adaptive inhibitors and the precision engineering of functional biomolecules.

## 2 Results

### 2.1 Overview of AnewSampling

AnewSampling is a transferable generative foundation framework designed for the high-fidelity sampling of all-atom equilibrium distributions of biomolecular complexes. To address the challenges of limited structural diversity in existing models, AnewSampling enables the faithful reproduction of MD accuracy across diverse systems, ranging from protein monomers to complex protein-ligand interactions. The framework is trained on AnewSampling-DB, the largest self-curated database of protein-ligand trajectories to date (Fig. 1c), which encompasses over 15 million conformations and 31,364 unique complexes.

The overall architecture of AnewSampling leverages a pretrained AlphaFold3-like architecture (Fig. 1b), integrating an Input Embedder, an MSA Module, and a Pairformer stack. To repurpose this architecture from deterministic static prediction to stochastic thermodynamic sampling, we implement a stratified hybrid fine-tuning strategy. Specifically, sequence representation modules are adapted via Low-Rank Adaptation (LoRA) to preserve their robust pretrained representation capabilities and mitigate the catastrophic forgetting of geometric reasoning. In contrast, the Diffusion Module undergoes full-parameter fine-tuning to accommodate the fundamental paradigm shift from deterministic structure prediction to a distributional sampler and to fully capture the complex manifold of distribution learning. A key aspect is the use of quotient-space diffusion [29]. (Fig. 1d). By mathematically factorizing out rigid-body degrees of freedom and constraining the generative process to the manifold of internal shapes, this formulation resolves the mathematical inconsistencies inherent in alignment-based training and ensures the precise recovery of the Boltzmann distribution.

To emulate the physical principle of ergodicity, we introduce a Cluster-Based Template Guidance mechanism (Fig. 1b). This approach decouples the target distribution from specific starting coordinates by randomly sampling and perturbing template structures from distinct conformational clusters, thereby enforcing an exhaustive exploration of the equilibrium ensemble from any valid initial state. Together, these advancements allow AnewSampling to investigate atomic interactions at an all-atom level while faithfully reproducing converged MD equilibrium distributions in a transferable manner.

### 2.2 Sampling Generalizes Across Unseen Conformational and Chemical Spaces

Current cutting-edge models reliably predict static structures but frequently fail to represent dynamic conformational distributions of protein-ligand complexes. To examine this systematically, we compared conformations generated by different models to a stringent ground-truth ensemble derived from MD simulations.

#### ATLAS Dataset

To comprehensively evaluate the capacity of AnewSampling to generate realistic conformational ensembles for protein monomers, we benchmarked our model on the ATLAS test set using standard metrics established in the AlphaFlow [33]. As detailed in Table 1, AnewSampling consistently outperforms existing state-of-the-art baseline methods—including BioMD [34], BioEmu [27]—across all major evaluation categories. Specifically, AnewSampling demonstrates unparalleled accuracy in predicting flexibility, achieving the highest Pearson correlations with reference REMD trajectories for Pairwise RMSD (*r* = 0.85) and pertarget RMSF (*r* = 0.93). Furthermore, our method exhibits exceptional distributional accuracy by minimizing the root mean Wasserstein-2 (W2) distance to 1.88 (compared to 2.18 for the next best method, BioMD) and successfully capturing the principal components of the MD conformational space. Crucially, AnewSampling excels at recovering transient ensemble observables, achieving state-of-the-art Jaccard indices for predicting weak contacts (*J* = 0.70) and highly transient contacts (*J* = 0.52). Collectively, these comprehensive metrics confirm that AnewSampling is a highly robust generative model capable of accurately reproducing the thermodynamic distributions and transient biophysical microstates inherent to monomeric protein dynamics.

**Table 1.** Evaluation of protein dynamics modeling on the ATLAS test set. Performance comparison of AnewSampling against state-of-the-art generative baselines for generating monomeric protein conformational ensembles. Performance metrics for all baseline methods evaluated on the ATLAS dataset are sourced from the BioMD literature. Arrows (↑ or ↓) adjacent to each metric designate whether higher or lower values indicate superior performance. Evaluated statistical measures include Pearson correlation (*r*), root mean Wasserstein-2 (W2) distance, Jaccard index (*J*), and Spearman correlation (*r*_*s*_). Bold text denotes the best performance achieved for each individual metric

| Category | Metrics | ESMFlow-MD (Full) | ESMFlow-MD (Distilled) | ConfDiff | BioEmu | Str2Str | MDGen | EBA | BioMD | AnewSampling |
| --- | --- | --- | --- | --- | --- | --- | --- | --- | --- | --- |
| Predicting flexibility | Pairwise RMSD $r(\uparrow)$ | 0.19 | 0.19 | 0.59 | 0.46 | 0.23 | 0.48 | 0.62 | 0.70 | **0.85** |
| | Global RMSF $r(\uparrow)$ | 0.31 | 0.33 | 0.67 | 0.57 | 0.38 | 0.50 | 0.71 | 0.76 | **0.89** |
| | Per-target RMSF $r(\uparrow)$ | 0.76 | 0.74 | 0.85 | 0.71 | 0.57 | 0.71 | 0.90 | 0.91 | **0.93** |
| Distributional accuracy | Root mean W2-dist. (↓) | 3.60 | 4.23 | 2.76 | 4.32 | 4.05 | 2.69 | 2.43 | 2.18 | **1.88** |
| | Trans. contrib. (↓) | 3.13 | 3.75 | 2.23 | 4.04 | 3.73 | - | 2.03 | 1.89 | **1.63** |
| | Var. contrib. (↓) | 1.74 | 1.90 | 1.40 | 1.77 | 1.43 | - | 1.20 | 1.10 | **0.89** |
| | MD PCA W2-dist. (↓) | 1.51 | 1.87 | 1.44 | 1.97 | 2.04 | 1.89 | 1.19 | 1.24 | **1.16** |
| | Joint PCA W2-dist. (↓) | 3.19 | 3.79 | 2.25 | 3.98 | 3.55 | - | 2.04 | 1.82 | **1.44** |
| | % PC-sim $> 0.5 \uparrow$ | 26 | 33 | 35 | **51** | 12 | - | 44 | 46 | **51** |
| Ensemble observables | Weak contacts $J(\uparrow)$ | 0.55 | 0.48 | 0.59 | 0.33 | 0.43 | 0.51 | 0.65 | 0.61 | **0.70** |
| | Transient contacts $J(\uparrow)$ | 0.34 | 0.30 | 0.36 | - | - | - | 0.41 | 0.46 | **0.52** |
| | Exposed residue $J(\uparrow)$ | 0.49 | 0.43 | 0.50 | - | - | 0.29 | 0.70 | 0.70 | **0.75** |
| | Exposed MI matrix $r_s(\uparrow)$ | 0.20 | 0.16 | 0.24 | 0.07 | 0.21 | - | 0.36 | 0.34 | **0.41** |

#### Held-out Test Set

We assessed the intrinsic flexibility of the ligands by calculating the Jensen-Shannon (JS) distance of ligand torsion angle distributions (Fig. 2a,b and Table S1). Models trained exclusively on static structures (AlphaFold3 (AF3) [6], Protenix (PX-v0.7.0) [8], Chai-1 [11]) exhibited median JS distances of approximately 0.5, indicating a profound inability to reproduce dynamic ligand distributions. Notably, although Boltz2 [10] incorporates limited MD data and B-factor during training, it similarly failed to capture these dynamics, yielding a median JS distance of 0.5297. This observation strongly suggests that training generative models via random structural perturbation of single, static conformations is fundamentally insufficient for learning the underlying physical energy landscape. In contrast, models trained on MD conformational data showed marked improvements. Strikingly, AnewSampling achieved a conformational distribution statistically indistinguishable from the MD baseline, yielding a median JS distance of 0.2402 (MD: 0.2251). Furthermore, AnewSampling maintained a consistently high success rate (JS distance ≤ 0.3) across diverse Pocket-SuCos similarity bins (a similarity partitioning metric adopted from IsoDDE [35]), demonstrating robust generalization to novel, out-of-distribution complexes (Fig. 2b).

**Figure 2.**
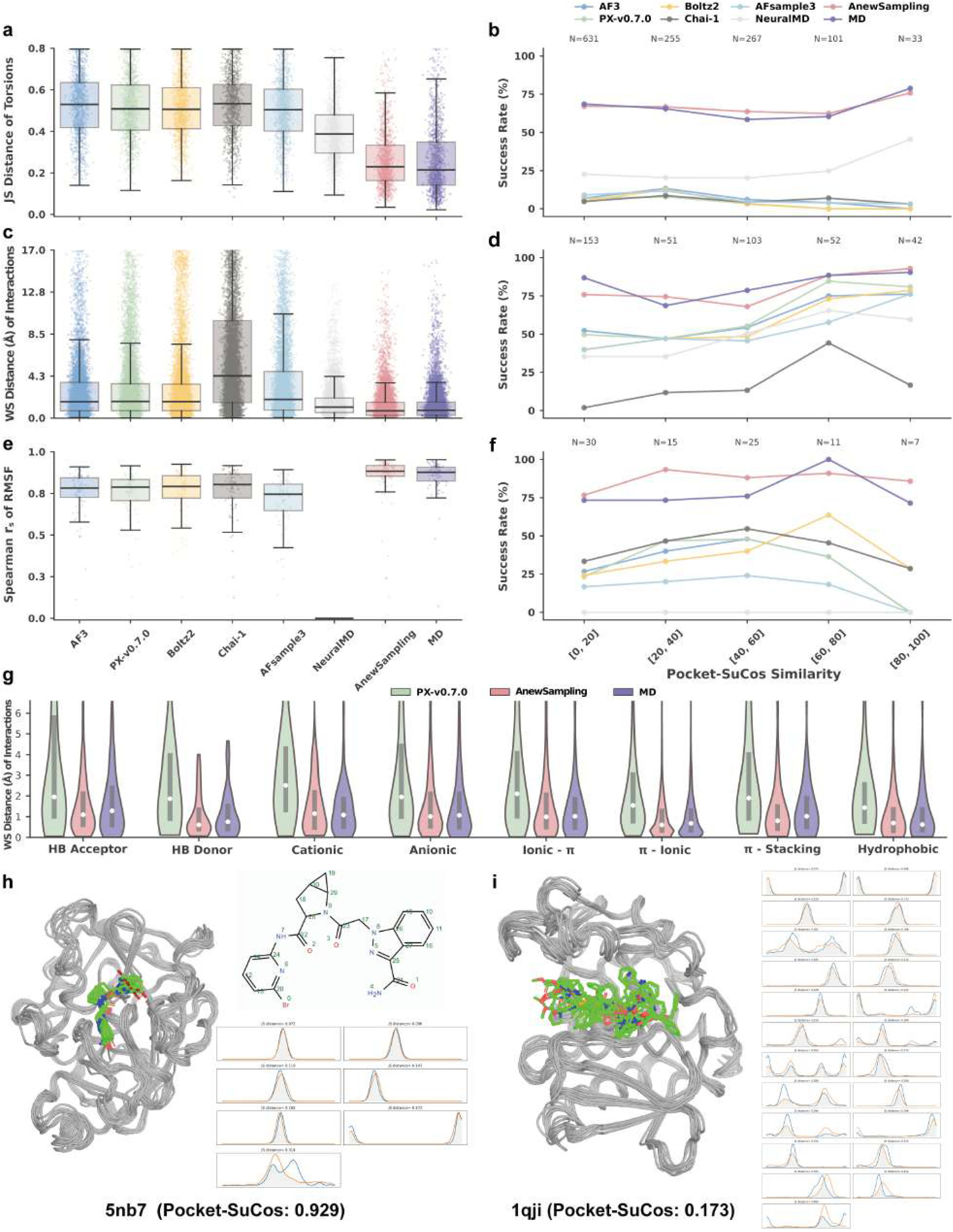
Evaluation of generative models for protein-ligand conformational ensemble generation on the held-out test set. **(a)** Boxplot of JS distances for ligand torsion angle distributions across 500 generated conformations compared to a 501-frame REMD reference ensemble. **(b)** Success rate (defined as JS distance ≤ 0.3) for ligand torsion predictions, stratified by Pocket-SuCos similarity to evaluate generalization capability. N indicates the number of torsions in each bin. **(c)** Boxplot of WS distances (in Å) comparing the distributions of protein-ligand interaction distances. This metric assesses non-covalent interaction stability without relying on reference-dependent superposition. **(d)** Success rate (defined as WS distance ≤ 0.3 Å) across different Pocket-SuCos similarity bins. N indicates the number of interactions in each bin. **(e)** Boxplot of Spearman correlation coefficients (*r*_*s*_) for protein backbone C*α* root-mean-square fluctuation (RMSF), evaluating the consistency of global protein dynamics. **(f)** Success rate (Spearman correlation ≥ 0.85) for capturing protein backbone dynamics across similarity bins. N indicates the number of systems in each bin. **(g)** Violin plots detailing the WS distance distributions for specific non-covalent interaction types (e.g., Hydrogen Bond Acceptor/Donor, Cationic, *π*-Stacking). **(h, i)** Structural overlays of generated protein backbones and ligand conformations, alongside corresponding ligand torsion probability density functions, for two representative test cases: a highly similar target (5nb7, similarity: 0.929) and a highly dissimilar target (1qji, similarity: 0.173).

Because ligand torsions do not fully capture the binding mechanism, we introduced a superposition-free metric using Wasserstein (WS) distance to evaluate the stability of specific atom-pair interactions within the binding pocket. While the co-folding models yielded median WS distances around 2.0 Å, AnewSampling accurately modeled the physical stability of these non-covalent interactions, achieving a median WS distance of 0.9931 Å—comparable to the MD baseline of 1.0561 Å (Fig. 2c and Table S2). For highly conserved interactions observed during MD simulations (frequency ≥ 0.8, Table S3), AnewSampling achieved a 66.3% success rate at a stringent WS distance threshold (≤ 0.3 Å), substantially outperforming the best co-folding model (AF3, 48.8%). This high-fidelity interaction modeling held true across all physical interaction types, including hydrogen bonds, ionic, and *π*-stacking interactions (Fig. 2g).

Beyond the binding pocket, we evaluated global protein dynamics by computing the Spearman correlation (*r*_*s*_) and root-mean-square error (RMSE) of the protein backbone C*α* RMSF (Fig. 2e and Table S4). AnewSampling effectively preserved global conformational motions, yielding a median RMSE of 0.2286, which surpassed both the best co-folding model (Boltz2: 0.7864) and the MD baseline (0.2699). Impressively, AnewSampling achieved a 75% success rate for highly correlated global dynamics (*r*_*s*_ ≥ 0.85), outperforming the MD reference (68%), and maintained this accuracy across all similarity bins (Fig. 2f).

#### In-house Dataset

To further validate the real-world applicability and robustness of AnewSampling, we extended our evaluation to an independent, internal dataset comprising 17 structurally diverse protein-ligand complexes (similarity: [0.06, 0.62]) derived from active drug discovery pipelines. Consistent with our test set benchmarks, AnewSampling maintained stable and high-fidelity conformational ensemble generation across these novel complexes. The model accurately reproduced local ligand flexibility, achieving a 68.8% success rate in torsion JS distance (≤ 0.3) (Fig S1b). It also successfully captured highly conserved protein-ligand interaction networks, yielding an 87.7% success rate (WS distance ≤ 0.3 Å) that closely trails the rigorous MD reference (97.3%) (Fig S1d). Furthermore, AnewSampling exhibited exceptional consistency in predicting global protein backbone dynamics, achieving a 100.0% success rate for highly correlated global motions (*r*_*s*_ ≥ 0.85) (Fig S1f), marginally outperforming the inherent variance of the MD baseline (94.1%). These findings underscore the capacity of AnewSampling to generalize effectively to novel, therapeutically relevant targets without degradation in thermodynamic sampling accuracy.

The transition from predicting static, single-state protein-ligand structures to generating thermodynamic ensembles represents a critical frontier in structure-based drug design. Our benchmarking clearly illustrates that while leading diffusion-based co-folding models (such as AF3 and Protenix) excel at finding local energy minima, they inherently lack the underlying physics required to sample a Boltzmann-like distribution of the target complex. AnewSampling overcomes this fundamental limitation by learning directly from MD-derived conformational spaces. The results unequivocally demonstrate that AnewSampling is the first all-atom generative model capable of producing highly accurate, equilibrium-state conformational distributions that mirror extensive REMD sampling (evidence in subsection 2.3). The model’s ability to maintain physical stability (evidenced by interaction WS distances) and global motion coherence (evidenced by RMSF correlation) across highly dissimilar targets highlights its robust generalization.

While AnewSampling achieves unprecedented parity with MD sampling for buried pocket interactions, our structural analysis (Fig. 2h,i) reveals minor deviations in the torsion distributions of highly solvent-exposed ligand moieties. We hypothesize that the current training paradigm captures the induced fit and pocket constraints well, but may lack sufficient sampling of unbound, solvent-driven ligand flexibility. Future iterations of AnewSampling could directly address this by incorporating large-scale, ligand-only MD datasets as a structural prior, thereby refining the generative distributions of solvent-facing functional groups and achieving true thermodynamic completeness.

### 2.3 Capturing perturbation-induced conformational shifts in SAR series

A fundamental challenge in lead optimization is that minor chemical modifications to a ligand—such as the addition of a methyl group, a halogen, or a subtle linker extension—can induce thermodynamically significant shifts in the conformational ensemble of the binding pocket. Conventional static structure prediction methods frequently neglect these subtleties, generally producing inflexible, fairly uniform pocket geometries over entire congeneric series. To investigate whether AnewSampling can detect and model these fine-grained structural responses, we evaluated our model on congeneric structure-activity relationship (SAR) series from the highly validated JACS and Merck benchmark datasets [31], comprising 593 complexes across 16 protein targets. Among these systems, where the median Pocket-SuCos similarity ranged from 0.53 to 0.93, our model consistently delivered outstanding performance. As shown in Fig. 3a. AnewSampling achieved the highest success rates among all evaluated generative methods across all three metrics, closely approaching the performance of the standard MD baseline.

**Figure 3.**
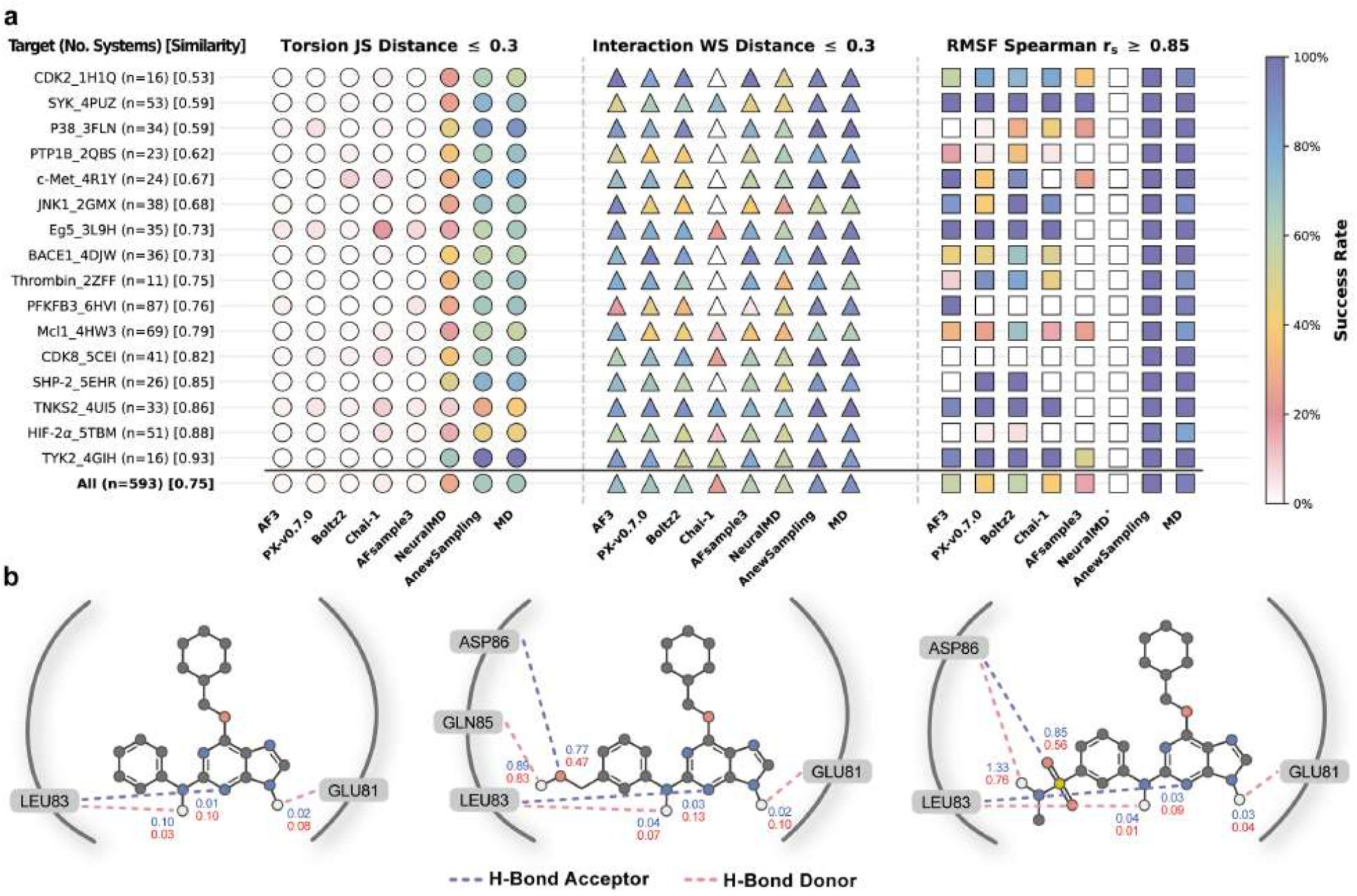
Evaluation of generative models for protein-ligand conformational ensemble generation on JACS and Merck datasets. **(a)** The heatmaps visualize the comparison between generative models and standard MD simulations in the success rates of ligand torsion JS distance ≤ 0.3 (left), protein-ligand highly conserved interaction (frequency ≥ 0.8) WS distance ≤ 0.3 Å (middle), and protein backbone C*α* RMSF Spearman correlation ≥ 0.85 (right). The color gradient reflects the success rate of each method, scaling from 0% (white) to 100% (dark blue). Each row corresponds to a specific protein target, arranged by its median pocket-SuCos similarity relative to the training set. Beside each target name and PDB ID, the number of evaluated ligands is indicated in parentheses, followed by the median Pocket-SuCos similarity in square brackets. ^*^Note that NeuralMD treats proteins as rigid stuctures, its corresponding RMSF values are uniformly zero. **(b)** Example of CDK2 ligands showing protein-ligand hydrogen bonds induced by phenyl ring substitutions and comparison in WS distances between MD (blue) and AnewSampling (red)

The success rate of torsion JS distance ≤ 0.3 specifically reflects how well a model reproduces the ligand conformational sampling distributions obtained from REMD (Fig. 3a left and Table S5). Static models—including AF3, Protenix, Boltz2, and Chai-1—inherently failed to reflect dynamic distributional shifts within the binding pocket. Consequently, they performed poorly on this metric, with maximum success rates below 20% and scoring below 2% for the vast majority of systems. While AFsample3 is designed to capture protein structural diversity, it similarly exhibited poor performance in predicting ligand conformational sampling. NeuralMD, which predicts trajectories for pocket-bound ligands, demonstrated marginal distribution prediction capabilities (overall success rate of 28%). In stark contrast, AnewSampling accurately captured these distributions, achieving an overall success rate of 66%, which is highly comparable to the sampling fidelity of standard MD.

Notably, standard MD simulations exhibited relatively low success rates in torsion JS distance for two specific targets: TNKS2 and HIF-2*α*. This discrepancy highlights inconsistant distributions sampled from standard MD and REMD, implying that these systems pose significant sampling challenges for conventional simulations. This difficulty stems from two primary factors. First, sampling specific torsions—such as the rotatable bonds between rings in TNKS2 ligands and the rotation of decorated phenyl rings in HIF-2*α* ligands—is hindered by high energy barriers. While MD frequently becomes kinetically trapped in local minima, REMD successfully traverses these barriers to sample multiple minima, resulting in divergent distributions. Second, some of the ligands in HIF-2*α* system may contain suboptimal initail local structures, such as incorrect sulfonyl

While fundamentally incapable of modeling dynamic ligand torsions, static co-folding models reproduced highly conserved protein-ligand interactions (frequency ≥ 0.8) reasonably well (Fig. 3a middle and Table S7), with AF3 performing the best among these methods. Although NeuralMD possesses some ligand conformational sampling capability, its interaction sampling performance was mediocre, likely bottlenecked by its rigid treatment of the protein. AnewSampling once again outperformed all other generative methods, achieving a remarkable success rate that closely mirrored the REMD baseline. This underscores AnewSampling’s proficiency in capturing subtle interaction shifts between congeneric ligands and the protein pocket. For instance, in the CDK2 system, which exhibited the lowest median Pocket-SuCos similarity (0.53), AnewSampling successfully captured the changes in pocket hydrogen bonding induced by phenyl ring substitutions while maintaining stable interactions in conserved regions (Fig. 3b). Furthermore, standard MD exhibited a high WS distance for the hydrogen bond between the sulfonamide group of molecule 28 and Asp, caused by insufficient sampling of both the ligand moiety and the adjacent pocket residues. AnewSampling’s ability to match the REMD distribution in this instance indicates a strong potential for enhanced sampling—a capability we discuss with examples in Section 2.4.

Beyond the accurate sampling of ligand torsions and pocket interactions, AnewSampling substantially surpassed other methods in predicting global protein dynamics, achieving the highest success rate for the protein C*α* RMSF Spearman correlation (*r*_*s*_ ≥ 0.85) across all systems (Fig. 3a right and Table S8). This demonstrates that AnewSampling maintains a high degree of biophysical realism in its global structural sampling. Finally, it is worth noting that because NeuralMD employs a strictly rigid protein representation, its calculated protein RMSF values are inherently zero, precluding it from capturing these dynamic fluctuations.

### 2.4 Enhanced Sampling Beyond Conventional MD

AnewSampling not only samples conformational distributions consistent with standard MD, but also explores regions that are difficult to reach within typical simulation budgets (approximately 1–100 ns). In such settings, enhanced-sampling methods such as REMD are often required to escape trapped local minima separated by high free-energy barriers. To demonstrate the capability of AnewSampling in navigating complex energy landscapes, we examined two ligands of the CDK2 receptor. CDK2 is a key member of the cyclin-dependent kinase family important in the regulation of the cell proliferation and RNA polymerase II transcription [32]. It has emerged as a critical target for tumor-selective therapy, particularly in CCNE1-amplified cancers, where its inhibition exploits synthetic lethality to eliminate tumor cells while sparing healthy tissues [36, 37]. Two of the protein ligand complexes for CDK2, 1h1r and 1h1s, are representative case necessitating REMD enhanced sampling to capture its multiple distinct binding poses [38]. As shown in the top penel of Figure 4, ligand 1h1r occupies two distinct binding poses characterized by a significant flip of the benzene ring, supported by the electron density of 1h1r in X-ray crystallography. While is challenging for standard MD escape the local minima of binding pose 1 of 1h1r, AnewSampling exhibits exceptional sampling capability by capturing both binding poses in agreement with the REMD simulations.

**Figure 4.**
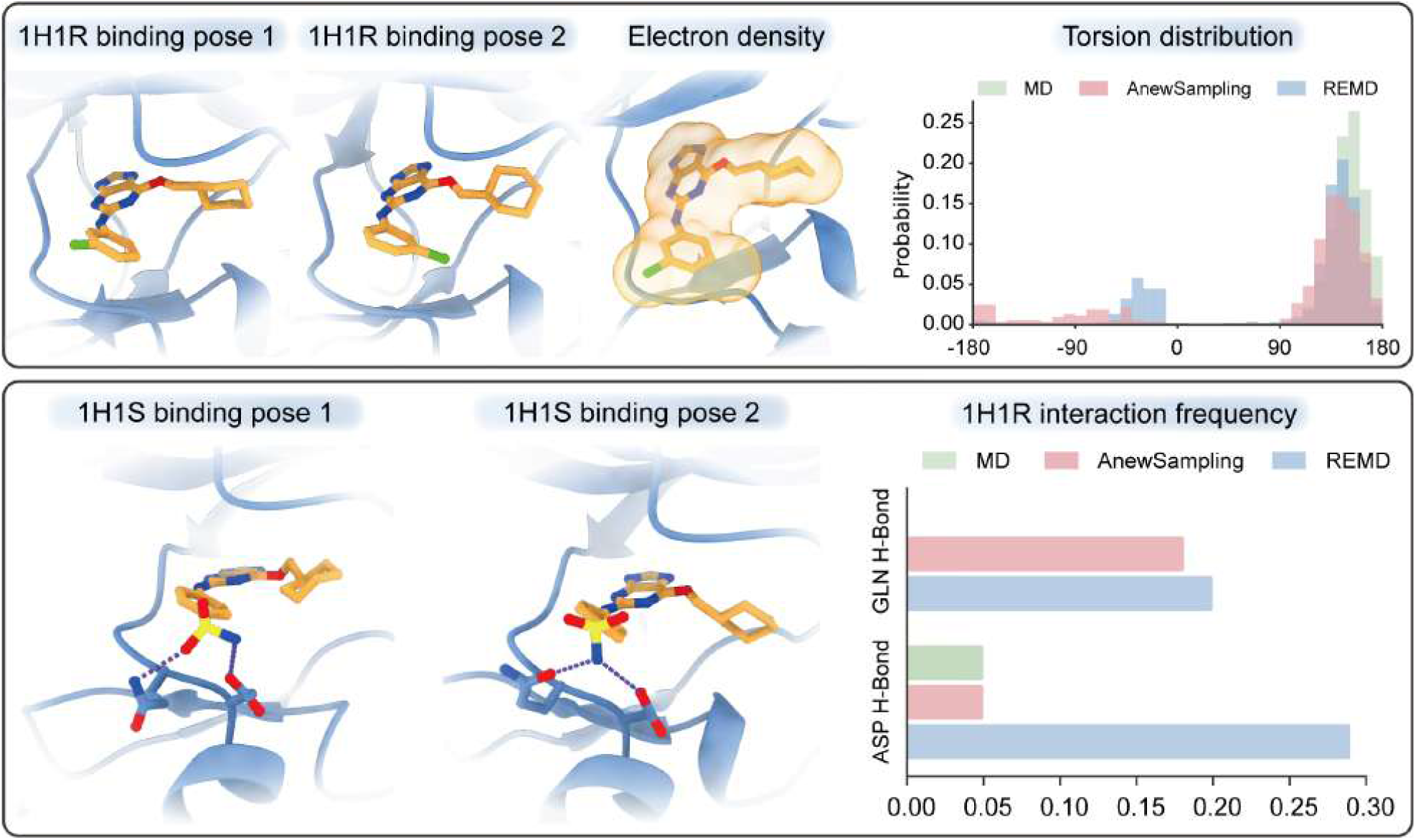
Enhanced conformational sampling in CDK2 complexes. (Top) The left panels illustrate two distinct binding poses of ligand 1h1r sampled from REMD and AnewSampling, alongside the experimental electron density of ligand 1h1r confirming the two alternative poses. The torsion distribution analysis (top right) demonstrates that conventional MD (green) is restricted to a single conformational basin (pose 1). In contrast, AnewSampling (pink) shows the potential of capturing the multimodal distribution observed in the REMD trajectory (blue). (Bottom) Visualization of two distinct 1h1s binding poses (left), highlighting the sulfonamide rotation and associated Gln/Asp side chain rearrangements required for hydrogen bond formation. Frequency analysis of the key interactions (right) confirms the ability of AnewSampling (pink) to capture the critical transient hydrogen bonds observed in REMD trajectory (blue), which are missed by conventional MD simulations (green).

Furthermore, we emphasize that accurate binding mode prediction often requires the coupled sampling of both ligand and protein degrees of freedom. This is exemplified by the ligand 1h1s as illustrated in the bottom panels of Figure 4, where the transition between binding modes involves not only the rotation of the ligand’s sulfonamide group but also the necessary reorganization of pocket residues (specifically Gln and Asp) to establish key hydrogen bond networks. Standard 10ns MD simulations typically fail to overcome the associated energy barriers, missing the critical Gln hydrogen bonds entirely. However, AnewSampling effectively captures these coupled side-chain fluctuations and recovers these transient hydrogen bonds with a fidelity comparable to REMD. This ability to accurately sample coupled protein-ligand fluctuations is vital for identifying bioactive conformations and improving the precision of activity predictions from both energetic and entropic perspectives. Although AnewSampling does not yet fully replicate the extensive configurational space exploration of REMD, it demonstrates remarkable enhanced sampling potential given that it was trained exclusively on standard MD data, which itself fails to capture these rare events. This suggests that the model effectively learns the underlying energy landscape rather than merely memorizing the limited conformational diversity of its training set.

## 3 Discussion

We have introduced AnewSampling, a generative foundation framework developed to sample the all-atom equilibrium distributions of biomolecular complexes. The framework has been demonstrated to achieve rigorous statistical alignment with ground-truth MD. And the generated ensembles yield evaluation metrics that are indistinguishable from the intrinsic variance observed between independent MD replicates. Statistically, the model establishes a new baseline by consistently outperforming all generative methods across every evaluation metric on the ATLAS monomer benchmark. Broadening the scope to protein-ligand dynamics, the framework accurately characterizes the precise atomic interactions. By capturing these fundamental interactions, the model faithfully recovers coupled ligand and side-chain motions within challenging targets such as CDK2 complexes, which are essential for industrial structure-activity relationship analysis. The computational inference operates at a fraction of the cost required for conventional MD simulations, effectively bypassing the high energy barriers that typically trap traditional sampling methods.

Despite the significant advancements introduced by AnewSampling, several limitations warrant further investigation in future research. Although the underlying architecture natively supports diverse biomolecular modalities, the primary constraint remains the scarcity of comprehensive training data. While we have curated the largest database to date, this database is insufficient to capture the full landscape of biological complexity. Future iterations will expand the framework to support a broader range of interaction types, such as protein-nucleic acid or protein-protein interactions. The current framework relies on the incorporation of templates. While this approach aligns with the standard inputs required for conventional MD, and the framework technically supports sequence-only inputs, the absence of templates results in diminished performance. The ultimate objective is to derive highly accurate equilibrium distributions solely from primary sequences. We observe that templates provide marginal benefits for simple monomeric systems; however, for complexes, the challenge becomes more pronounced. As observed in models similar to AlphaFold 3, accurately predicting the static structures of such complexes is inherently difficult. Extending this to the task of distribution prediction further exacerbates the challenge. Consequently, scaling the training data and refining the architectural structures represent critical directions for future work.

The current framework learns thermodynamic distributions within a single, fixed thermodynamic environment. In contrast, conventional MD can be naturally extended to model varying macroscopic conditions, although it still encounter severe sampling bottlenecks. Our AnewSampling framework and conventional MD are fundamentally complementary. MD provides the essential raw training data that underpins the generative system. As previously noted, by learning fundamental physical interactions across diverse chemical systems, AnewSampling demonstrates an emergent capacity for enhanced sampling. Conversely, the generative framework can accelerate MD by providing a diverse and robust set of initial structural candidates. By utilizing these generated conformations as starting points, traditional physical simulations can efficiently bypass potential energy barriers and explore the broader conformational space with unprecedented thoroughness.

## 4 Method

### 4.1 Datasets

We constructed the AnewSampling-DB dataset, which represents a significant advancement in both scale and physical fidelity, by filtering and curating data from PDBBind v2020 [39], ChEMBL [40], JACS and Merck [31]. The dataset comprises 31,364 complexes, substantially outperforming the MISATO dataset [41] (15,592 complexes) while covering a broad chemical space of 10,297 unique protein sequences and 27,979 unique ligand SMILES. Building upon the logic of the Bindingnet v2 hierarchical template-based modeling approach [42], we employed Glide [43] to prepare ~20,000 complexes from ChEMBL [40] to enrich our dataset with congeneric series. The inclusion of congeneric series captures the subtle differences in protein-ligand interactions between structurally similar ligands with distinct bioactivities binding to identical protein targets. The dataset exhibits robust distributional characteristics across key structural metrics. The torsion number distribution (Fig. 5B) shows a unimodal peak between 4 and 6 rotatable bonds while extending to highly flexible ligands, encapsulating over 43,000 distinct force field torsion types. Token and sequence number distributions further highlight the dataset’s coverage of diverse system sizes.

**Figure 5.**
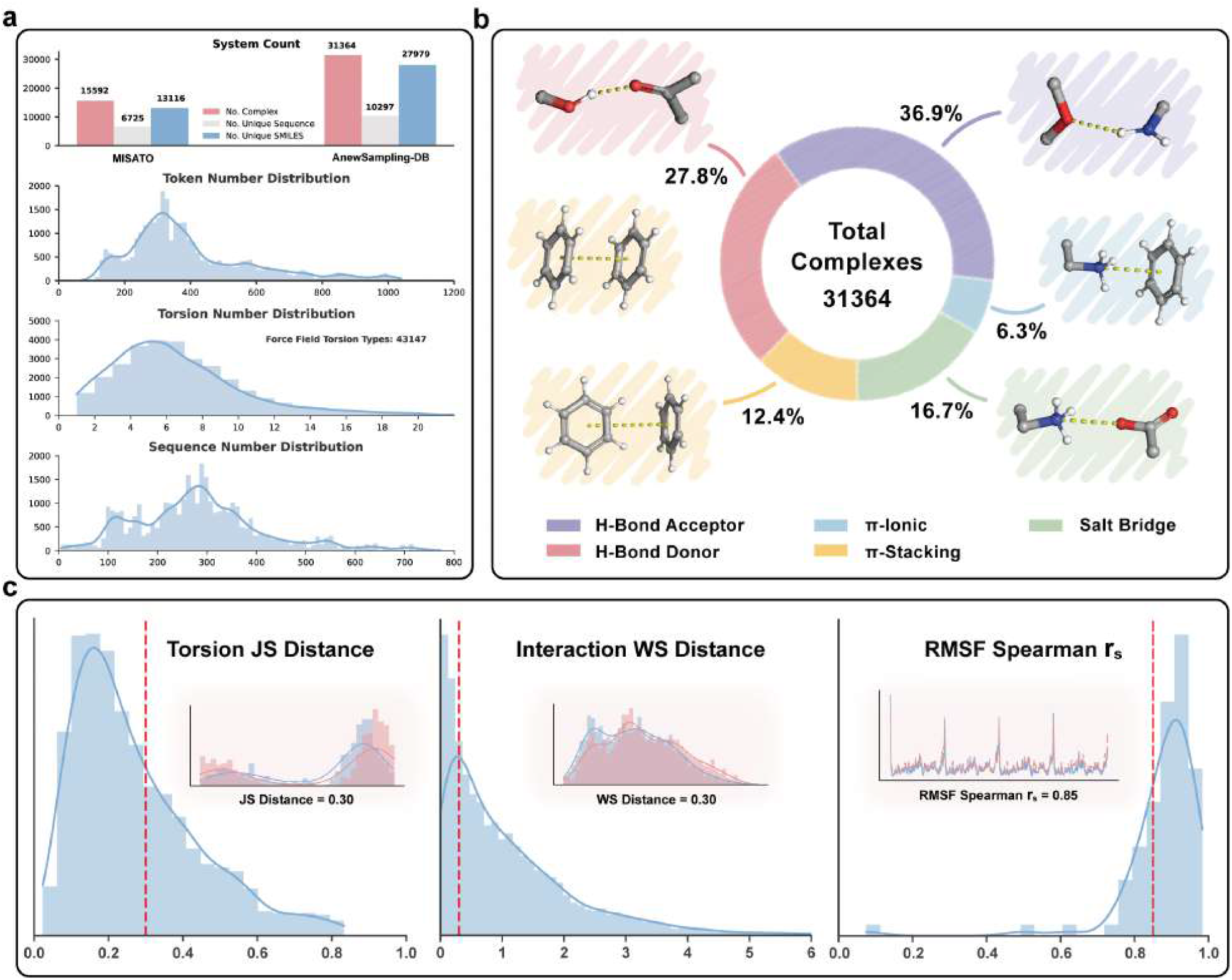
Statistical overview of the AnewSampling-DB dataset and Evaluation Metrics. **(a)** Classification of polar and aromatic interaction types within the 31,364 complexes. The percentage breakdown details specific contacts (e.g., H-bonds, *π*-stacking), excluding hydrophobic and Van der Waals interactions which constitute the majority (67.7%) of the total profile. **(b)** Dataset scale comparison against the MISATO benchmark, highlighting the increase in unique protein sequences and ligand SMILES. Subplots display the distributions of token counts, torsion numbers (covering 43147 types), and sequence lengths. **(c)** Examples showing the evaluation metrics of torsional JS distance (JSD), interaction WS distance, and protein C*α* RMSF.

To capture the complexity of molecular recognition, the dataset balances a wide array of non-covalent interaction types. The overall interaction landscape is dominated by hydrophobic and van der Waals (vdW) contacts, which collectively account for 67.7% of the total interactions. Focusing on the specific polar and aromatic interactions (excluding hydrophobic and vdW contacts), Fig. 5A illustrates a distribution primarily driven by hydrogen bonding, where H-bond acceptors and donors constitute 36.9% and 27.8% of this subset, respectively. This specific interaction profile is further enriched by essential electrostatic and aromatic contacts, including salt bridges (16.7%), *π*-stacking (12.4%), and *π*-ionic interactions (6.3%).

To ensure high data quality and physical consistency across this expansive library, all molecular dynamics (MD) trajectories were generated via a unified pipeline employing a consistent set of protein and ligand force fields. Furthermore, REMD simulations were utilized to rigorously explore the conformational landscape more efficiently. Detailed simulation protocols are provided in the Appendix.

### 4.2 Evaluation Metrics

To systematically quantify generative performance, we established strict success thresholds grounded in physical intuition and empirical reference distributions (Fig. 5c). For intrinsic ligand flexibility in the binding pocket, we defined successful recovery as a JS distance of ≤ 0.30 for torsion angle distributions, a threshold representing highly overlapping conformational states. To evaluate the precise stability of protein-ligand contact networks without relying on rigid-body superposition, we set a stringent WS distance threshold of ≤ 0.30 Å. Finally, to ensure the preservation of coherent global protein backbone dynamics, we required a Spearman correlation (*r*_*s*_) of ≥ 0.85 for the RMSF. These rigorously defined metrics ensure that generated ensembles are evaluated not just on static geometric similarity, but on true thermodynamic and dynamic fidelity.

## Appendix

### A Methods

#### A.1 Evaluation Metrics

##### ATLAS Evaluation Metircs

Following the benchmarking frameworks established by AlphaFlow [33] and BioMD [34], we evaluated AnewSampling using the ATLAS dataset [30] to perform ensemble analysis (predicting flexibility, distributional accuracy, and ensemble observables). Prior to analysis, all ensembles were aligned to the initial static all-atom structure. Computations were performed using Cartesian coordinates via MDTraj [44], with trajectories subsampled to 1,000 frames to ensure computational efficiency. The 2-Wasserstein distance between two 3D Gaussian distributions is defined as:

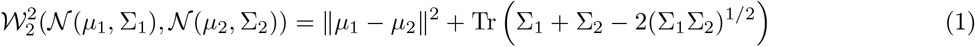

Notably, this metric generalizes the Euclidean distance metric used for point masses in AlphaFlow. The aggregate Root Mean Squared Wasserstein Distance (RMWD) can thus be decomposed into distinct translational and variance components:

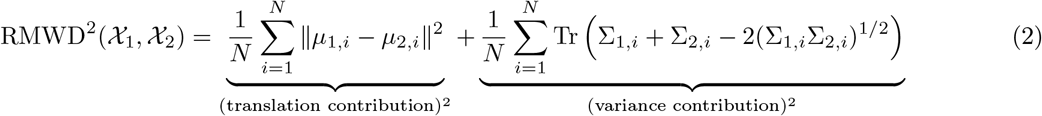

Quantitative results for these contributions are detailed in Table S1. For joint _2_ distance calculations, we projected the data into a PCA subspace to mitigate convergence issues arising from high-dimensional thermal fluctuations. To avoid biasing the projection solely towards the reference simulation, which may obscure orthogonal deviations in the predicted ensemble, we performed PCA using both the MD reference ensemble alone and a pooled ensemble (equal weighting of MD and predicted frames). Finally, sidechain solvent-accessible surface area (SASA) was computed using the Shrake-Rupley algorithm (probe radius: 2.8 Å), with residues classified as buried if their SASA fell below 2.0 Å^2^ [45].

##### Ligand Conformational Fidelity via Torsional JS Distance

To assess how accurately our model explores the internal degrees of freedom of the ligand within the binding pocket, we utilize the torsional JS distance (*D*_*JS*_). We compute the probability distribution of all rotatable dihedrals (*ϕ*) for the ligand across the generated ensemble *P* and the MD reference ensemble *Q*:

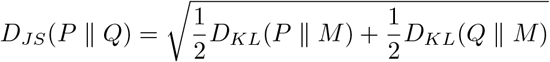

where 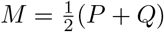. This metric quantifies the divergence between predicted and ground-truth torsional preferences, ensuring that the model captures physical rotameric states rather than merely predicting averaged geometries. For each rotatable bond, we discretize the dihedral angle space into *N* = 36 bins (bin size = 10°) to construct a discrete probability distribution of observed orientations.

##### Interaction Distance Distribution Difference via WS Distance

The stability of a protein-ligand complex is governed by its interaction network (e.g., hydrogen bonds, *π*-stacking, and salt bridges). To evaluate the sampling of these non-bonded interactions, we characterized the dynamic protein-ligand interaction landscape using the ProLIF toolkit [46] across 500 frames. We calculate the WS distance (*W*_1_) between the distributions of specific protein-ligand atomic distances:

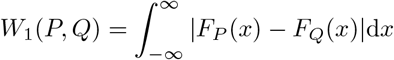

The *W*_1_ metric (or “Earth Mover’s Distance”) provides a continuous measure of how much “work” is required to transform the predicted interaction profile into the MD-derived profile. Crucially, the calculation accounts for the inherent symmetry of ligands, residues, and protein chains; equivalent atoms and subunits are permuted to minimize the distance, thereby preventing structural degeneracies from inflating the error metric. By integrating across the entire ensemble, this metric penalizes both the omission of critical binding modes and the generation of energetically unfavorable clashes.

##### Global Protein Dynamics via C*α* RMSF

To determine if the generative model recovers the intrinsic plasticity of the protein scaffold, we compare the C*α* RMSF across multiple diverse systems.

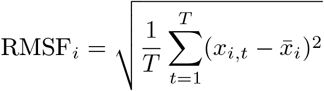

By correlating the predicted RMSF profiles with MD benchmarks across a broad test set, we demonstrate the model’s ability to distinguish between rigid secondary structures and disordered loops. This global assessment ensures that the ligand-binding event is coupled with realistic protein transitions, maintaining the biophysical integrity of the holo-structure.

#### A.2 Simulation Protocol

The GROMACS 2023 package [47] was used to conduct the all-atom MD simulations. The ligand-protein complexes were prepared by placing the molecules at the center of triclinic boxes with a minimum distance of 1.0 nm to the box boundary. The TIP3P water model [48] was used to solvate the complexes. The system was neutralized and brought to a physiological ionic strength of 0.15 M by adding sodium and chloride ions [49]. The proteins were parameterized with Amber ff14SB [50] and ligands were parameterized with the ByteFF [51]. The relaxation of each system started with energy minimization of 50,000 steps, followed by 400 ps relaxation in the *NV T* ensemble with a 1fs time step to increase the temperature to 298.15K and 100 ps relaxation in the *NPT* ensemble at 298.15 K and a pressure of 1 bar. The protein backbone atoms and the ligand heavy atoms were restrained by harmonic position restraints with a force constant of 1000 kJ mol^−1^nm^−2^ during relaxation. Finally, the systems were further equilibrated without any restraints for 2 ns in the *NPT* ensemble with a 2 fs time step before collecting 10 ns production data with saving trajectory every 20 ps. For the REMD simulations, we adopted the replica exchange with solute tempering (REST2) [52] enhanced sampling scheme with all the ligand atoms selected as the hot region. A total of 16 *λ* windows were used for each REMD simulation to scale the nonbonded and torsional interactions involving the hot region. The alchemical schedule of effective temperature of the hot region was estimated with the expected acceptance ratio of replica exchange set to be 0.3 [53].

The temperature in the simulations was controlled using a Langevin integrator [54] with the friction coefficient set to 0.5 ps^−1^. The stochastic cell rescaling algorithm [55] was applied for pressure coupling with a time constant of 2 ps. All the bonds with hydrogen atoms were constrained using the LINCS algorithm [56]. Periodic boundary conditions were applied, and the particle mesh-Ewald method [57] was used to treat long-range electrostatic interactions with a direct space cutoff of 1.0 nm. The Lennard-Jones interactions were switched off at 0.8 nm and a dispersion correction was applied for energy and pressure. The analysis of MD trajectories was performed using MDAnalysis [58, 59] and ProLIF [46].

#### A.3 Structural Refining Protocol

Our structural refining protocol emphasized the preservation of biological and chemical integrity. We employed PDBFixer [60] and Schrödinger Prime [61] to model missing loops and non-standard amino acids. To preserve the structural integrity of protein chains, we conducted global alignments between the sequences derived from the PDB and their respective RefSeq identities. Protein-ligand complexes were prepared using Schrödinger protein preparation wizard [62]. Protonation states were generated with Epik [63] then optimized with PROPKA [64].

#### A.4 Quantification of system similarity

We used a combined similarity measure to carefully test how well our model could generalize to different, out-of-distribution targets. This tool measures the distance between different protein-ligand complexes by taking into account both ligand-and pocket-centric factors. It is based on the stratification method set up in the IsoDDE report [35]. This particular metric, called Pocket-SuCos [65], blends a ligand-focused SuCos score—which only looks at three-dimensional volumetric overlap and physicochemical feature similarity (feature score)—with a pocket query coverage (pocket_qcov) score to make sure that the binding site stays the same.

#### A.5 Baselines and Implementation Details

All benchmark evaluations were conducted on a local high-performance computing cluster using a standardized hardware baseline to ensure consistency in the inference environment. To ensure a rigorous and fair comparison, each baseline model was deployed using its official local repository, with input protocols optimized to match the information density of AnewSampling:

- **Protenix (PX-v0.7.0)**: We employed the official base version of Protenix. Similar to other co-folding models, the inference script was adapted to accept 3D ligand conformations (SDF) as spatial hints, ensuring the model had access to the same initial geometric information as AnewSampling.
- **Boltz2**: The official Boltz2 open-source codebase was used for local inference. The input protocol was modified to accept SDF conformations. During evaluation on the held-out test set, 1 system failed due to Out-of-Memory (OOM) errors and was subsequently excluded from the aggregate statistics.
- **Chai-1**: We utilized the official Chai-1 repository. The input logic was similarly modified to support 3D SDF conformations. Due to its substantial memory footprint when processing complex targets in the held-out test set, 9 systems encountered OOM errors; these cases were omitted from the final performance metrics.
- **NeuralMD**: This model was initialized with the starting frame of each system and performed continuous sampling for 500 steps. It serves as a dedicated baseline for evaluating the propagation of local molecular dynamics from a known starting point.

##### Sampling Strategy and Ensemble Construction

To ensure a direct comparison with the ensemble generation capabilities of AnewSampling, we applied a uniform enhanced sampling protocol to all co-folding baselines (Protenix, Boltz2, and Chai-1). For each case in the test set, we executed the models using 5 distinct random seeds, with each seed generating 100 independent conformations. The resulting 500 frames were aggregated to construct the structural ensemble for each model, which was then used to calculate distribution-based metrics, including JS distance of ligand torsions, WS distance of interaction pairs, and RMSF of protein C*α*.

**Table S1.**
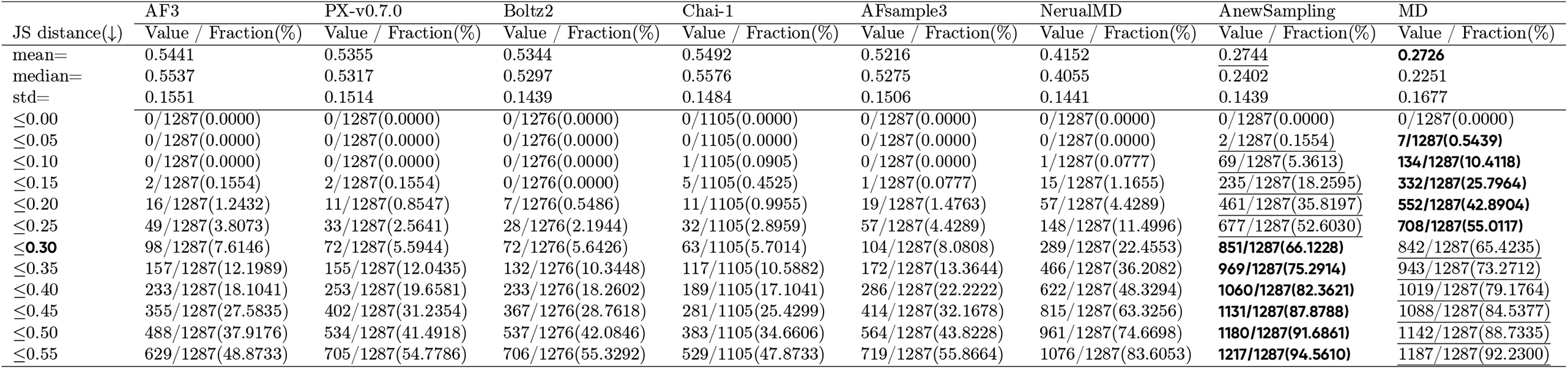
Statistical summary of JS distances for ligand torsion angle distributions on the held-out testset. The table provides the mean, median, standard deviation, and cumulative success rates (binned from ≤ 0.00 to ≤ 0.55) of JS distances for conformations generated by the co-folding models, MD-trained generative models, and the MD baseline.

**Table S2.**
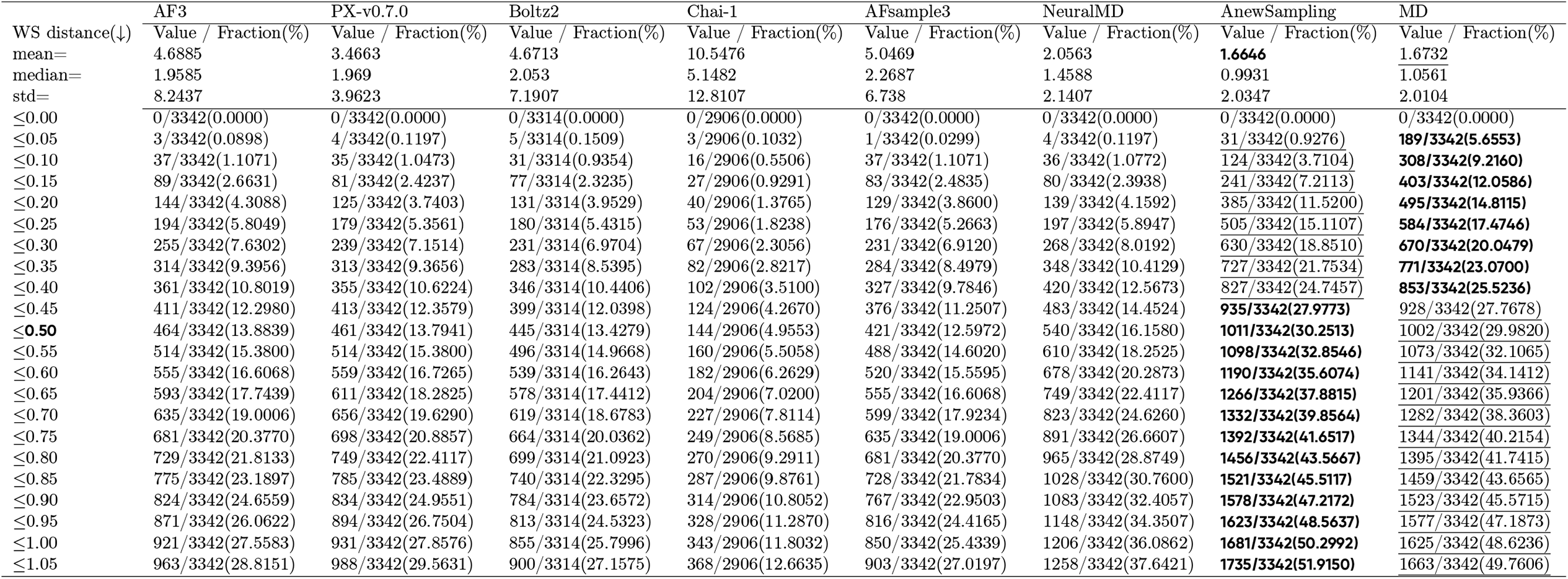
Wasserstein distance statistics for all overall protein-ligand interactions on the held-out testset. Data includes mean, median, standard deviation, and binned cumulative frequencies for WS distances mapping the continuous stability of non-covalent interactions within the binding pocket

**Table S3.**
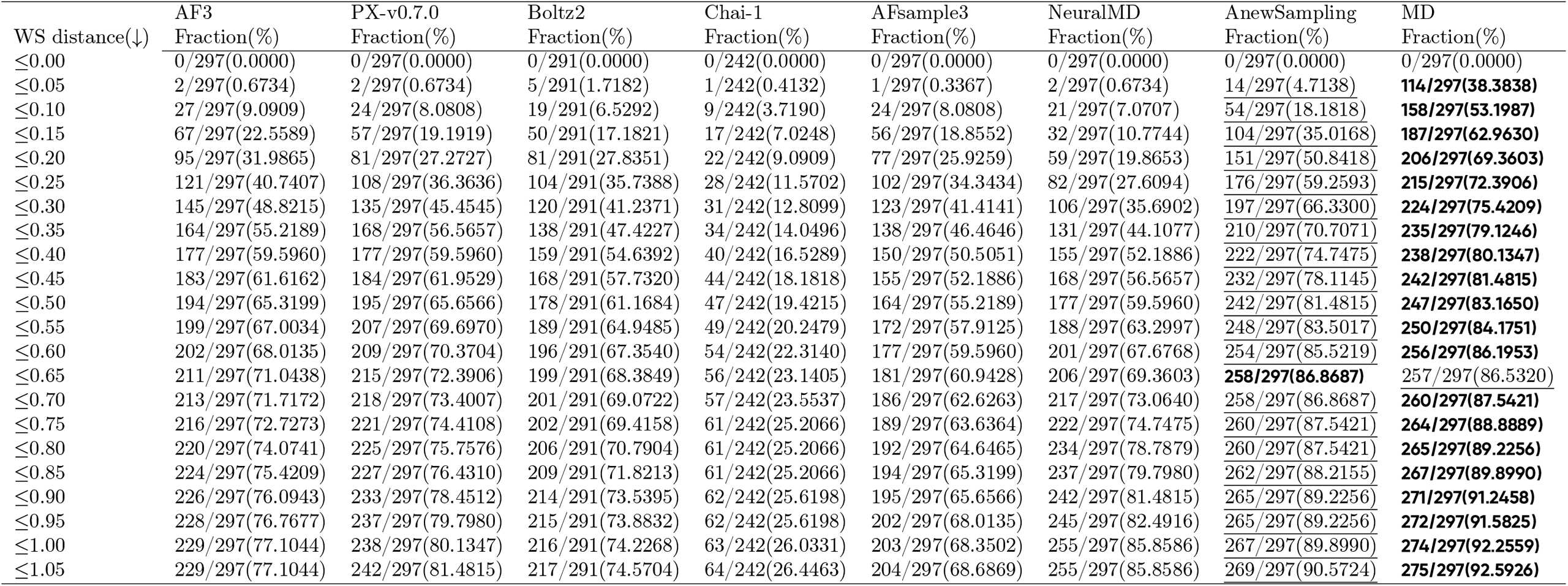
Wasserstein distance statistics for highly conserved protein-ligand interactions on the held-out testset. This table isolates structural interactions that occur with high frequency (≥ 0.8) in the reference REMD trajectories, highlighting the capacity of each model to capture strongly bound, dominant interaction states.

**Table S4.**
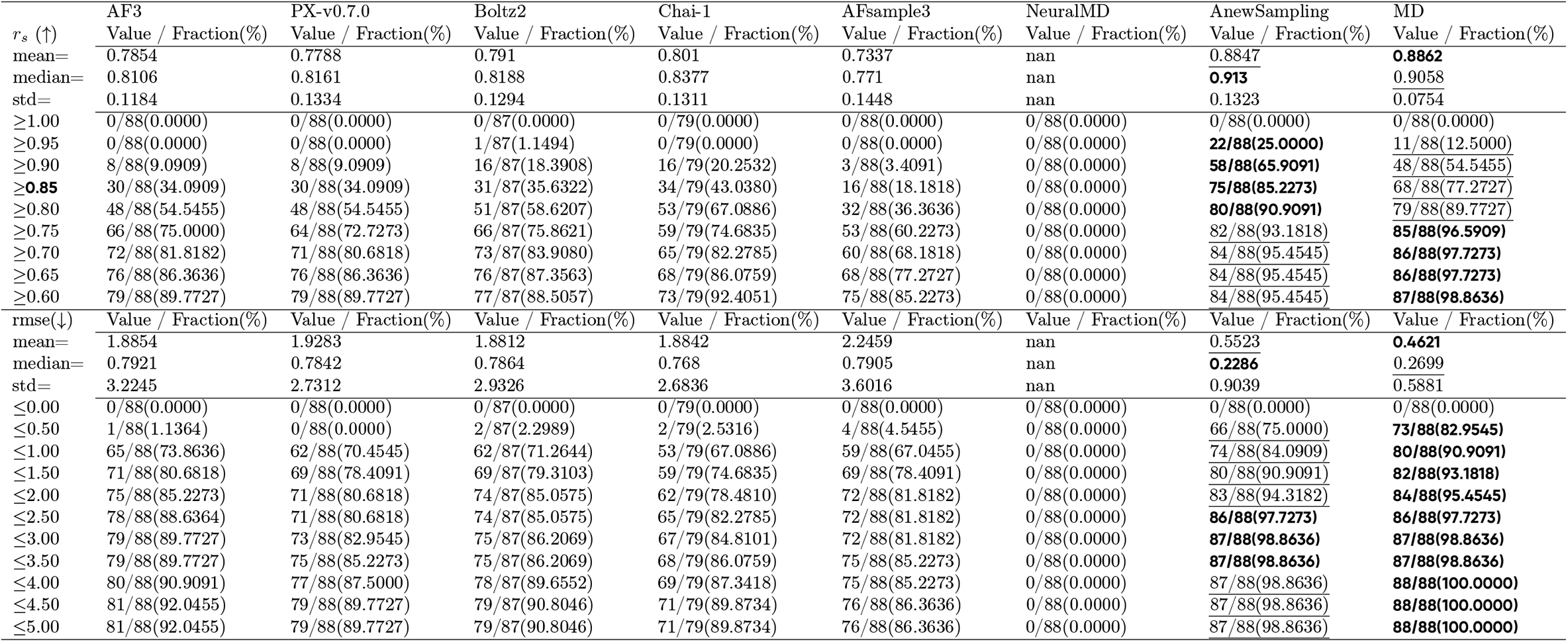
Evaluation metrics for protein backbone structural dynamics on the held-out testset. Comparative performance of each model in predicting global structural flexibility, quantified via Spearman correlation *r*_*s*_ and RMSE of the protein backbone C*α* RMSF relative to the REMD ground truth

**Figure S1.**
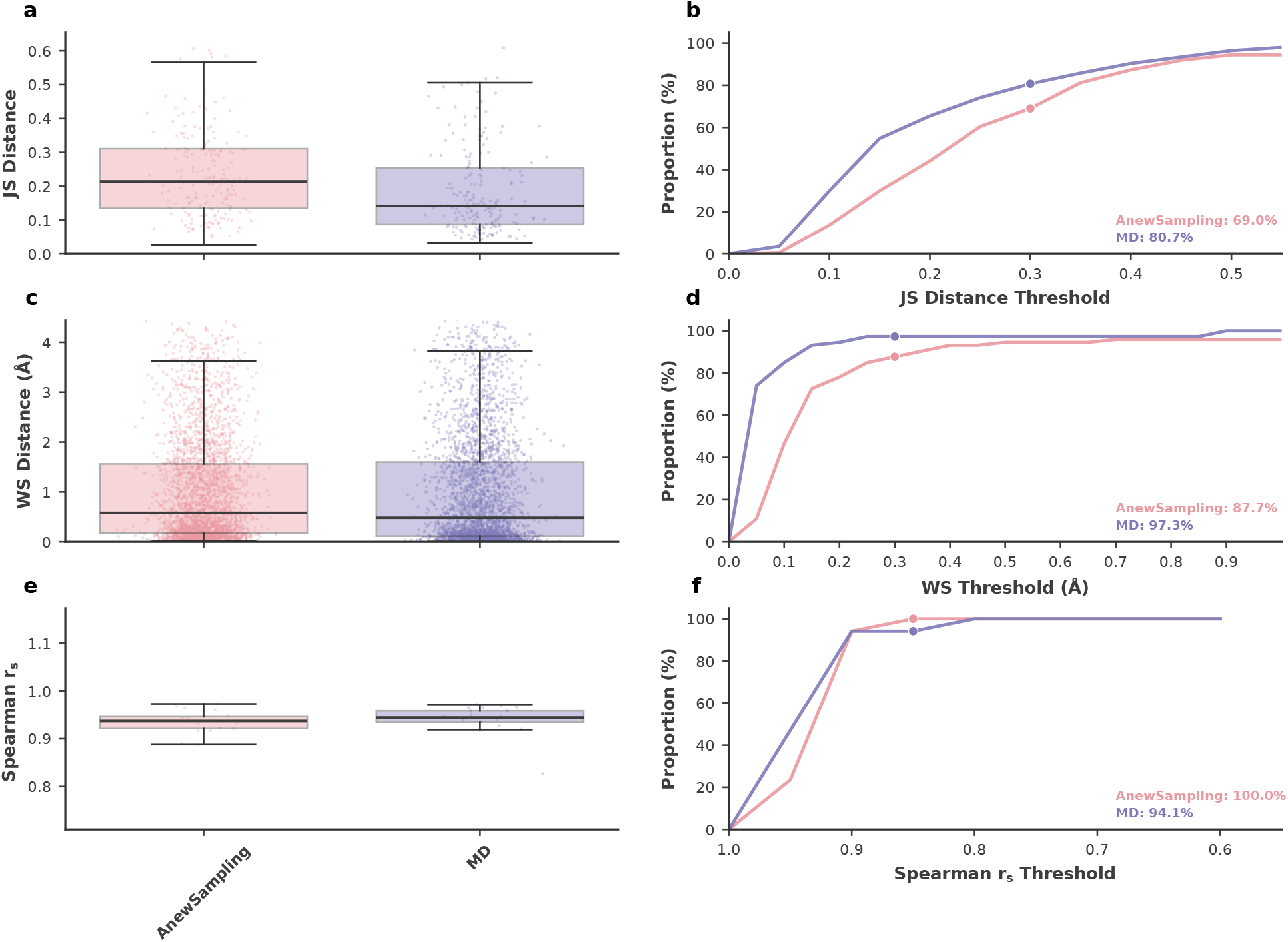
Evaluation of generative models for protein-ligand conformational ensemble on an independent internal pipeline dataset. **(a)** Boxplot comparing the JS distance of ligand torsion angle distributions between AnewSampling and the MD baseline. **(b)** Cumulative success rate of torsion JS distances; AnewSampling achieves a 68.8% success rate at a threshold of ≤ 0.3, compared to 80.2% for the MD baseline.**(c)** Boxplot detailing the WS distance (in Å) for protein-ligand interaction distributions. **(d)** Cumulative success rate for highly conserved, stable interactions (MD frequency ≥ 0.8), with AnewSampling reaching an 87.7% success rate at a stringent WS threshold of ≤ 0.3 Å. **(e)** Boxplot of RMSE for protein backbone C*α* RMSF, demonstrating the consistent prediction of global protein flexibility. **(f)** Cumulative success rate of *r*_*s*_ for protein RMSF; AnewSampling exhibits a 100.0% success rate at *r*_*s*_ ≥ 0.85, demonstrating robust recovery of coherent structural dynamics across the internal dataset.

**Table S5.** Statistical summary of JS distances for ligand torsion angle distributions on the jacsmerck dataset. The table provides the mean, median, standard deviation, and cumulative success rates (binned from ≤ 0.00 to ≤ 0.55) of JS distances for conformations generated by the co-folding models, MD-trained generative models, and the MD baseline.

|  | AF3 | PX-v0.7.0 | Boltz2 | Chai-1 | AFsample3 | NeuralMD | AnewSampling | MD |
| --- | --- | --- | --- | --- | --- | --- | --- | --- |
| JS distance(↓) | Value/Fraction(%) | Value/Fraction(%) | Value/Fraction(%) | Value/Fraction(%) | Value/Fraction(%) | Value/Fraction(%) | Value/Fraction(%) | Value/Fraction(%) |
| mean= | 0.5844 | 0.5910 | 0.5869 | 0.5589 | 0.5653 | 0.3854 | 0.2659 | 0.2640 |
| median= | 0.5918 | 0.5937 | 0.5934 | 0.5676 | 0.5753 | 0.3720 | 0.2395 | 0.2251 |
| std= | 0.1165 | 0.1199 | 0.1217 | 0.1321 | 0.1208 | 0.1329 | 0.1421 | 0.1562 |
| $\leq 0.00$ | 0/3060(0.0000) | 0/3060(0.0000) | 0/3060(0.0000) | 0/3060(0.0000) | 0/3060(0.0000) | 0/3060(0.0000) | 0/3060(0.0000) | 0/3060(0.0000) |
| $\leq 0.05$ | 0/3060(0.0000) | 0/3060(0.0000) | 0/3060(0.0000) | 0/3060(0.0000) | 0/3060(0.0000) | 0/3060(0.0000) | 12/3060(0.3922) | 23/3060(0.7516) |
| $\leq 0.10$ | 0/3060(0.0000) | 0/3060(0.0000) | 0/3060(0.0000) | 0/3060(0.0000) | 0/3060(0.0000) | 4/3060(0.1307) | 251/3060(8.2026) | 335/3060(10.9477) |
| $\leq 0.15$ | 0/3060(0.0000) | 0/3060(0.0000) | 1/3060(0.0327) | 2/3060(0.0654) | 0/3060(0.0000) | 48/3060(1.5686) | 697/3060(22.7778) | 842/3060(27.5163) |
| $\leq 0.20$ | 2/3060(0.0654) | 5/3060(0.1634) | 8/3060(0.2614) | 18/3060(0.5882) | 3/3060(0.0980) | 191/3060(6.2418) | 1195/3060(39.0523) | 1328/3060(43.3987) |
| $\leq 0.25$ | 10/3060(0.3268) | 12/3060(0.3922) | 17/3060(0.5556) | 53/3060(1.7320) | 14/3060(0.4575) | 465/3060(15.1961) | 1614/3060(52.7451) | 1706/3060(55.7516) |
| $\leq 0.30$ | 36/3060(1.1765) | 39/3060(1.2745) | 42/3060(1.3725) | 114/3060(3.7255) | 57/3060(1.8627) | 872/3060(28.4967) | 2027/3060(66.2418) | 2037/3060(66.5686) |
| $\leq 0.35$ | 101/3060(3.3007) | 89/3060(2.9085) | 105/3060(3.4314) | 225/3060(7.3529) | 145/3060(4.7386) | 1302/3060(42.5490) | 2314/3060(75.6209) | 2269/3060(74.1503) |
| $\leq 0.40$ | 204/3060(6.6667) | 188/3060(6.1438) | 218/3060(7.1242) | 373/3060(12.1895) | 314/3060(10.2614) | 1788/3060(58.4314) | 2540/3060(83.0065) | 2474/3060(80.8497) |
| $\leq 0.45$ | 434/3060(14.1830) | 366/3060(11.9608) | 417/3060(13.6275) | 622/3060(20.3268) | 543/3060(17.7451) | 2173/3060(71.0131) | 2697/3060(88.1373) | 2644/3060(86.4052) |
| $\leq 0.50$ | 685/3060(22.3856) | 672/3060(21.9608) | 701/3060(22.9085) | 951/3060(31.0784) | 882/3060(28.8235) | 2492/3060(81.4379) | 2820/3060(92.1569) | 2768/3060(90.4575) |
| $\leq 0.55$ | 1088/3060(35.5556) | 1082/3060(35.3595) | 1104/3060(36.0784) | 1393/3060(45.5229) | 1288/3060(42.0915) | 2710/3060(88.5621) | 2925/3060(95.5882) | 2874/3060(93.9216) |

**Table S6.**
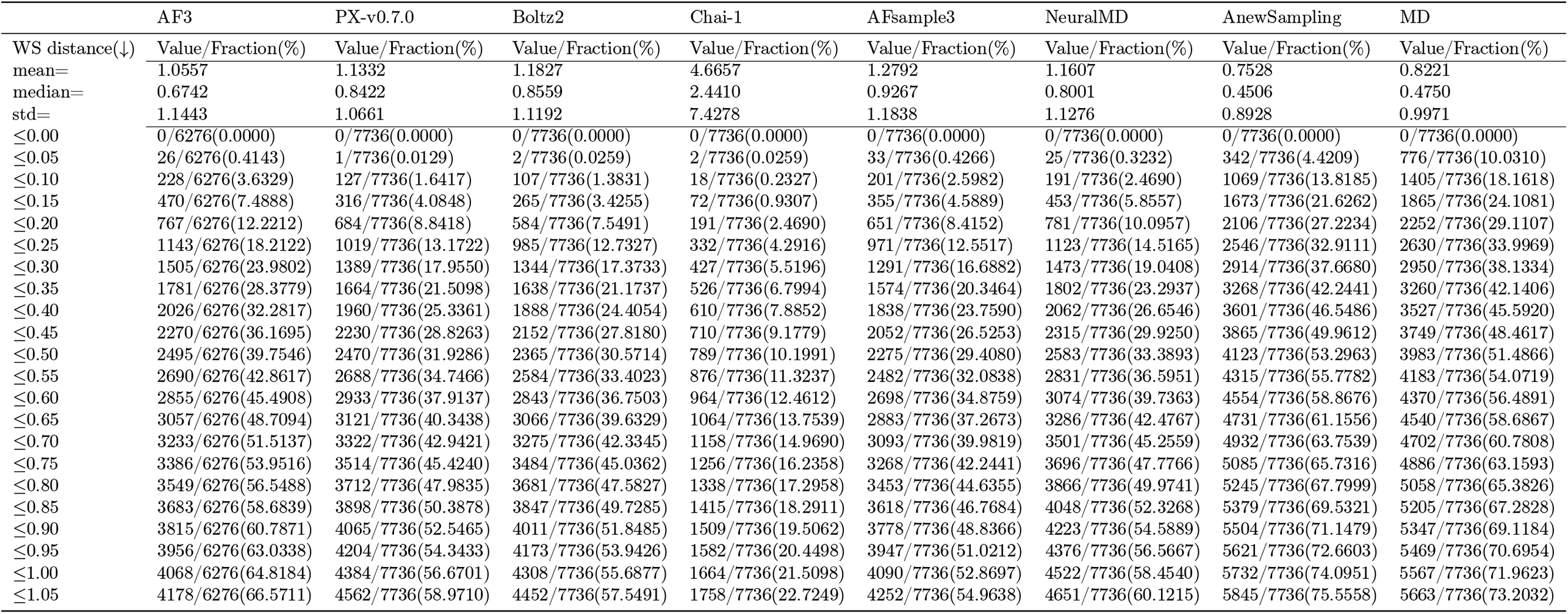
Wasserstein distance statistics for all overall protein-ligand interactions on the jacsmerck dataset. Data includes mean, median, standard deviation, and binned cumulative frequencies for WS distances mapping the continuous stability of non-covalent interactions within the binding pocket.

**Table S7.**
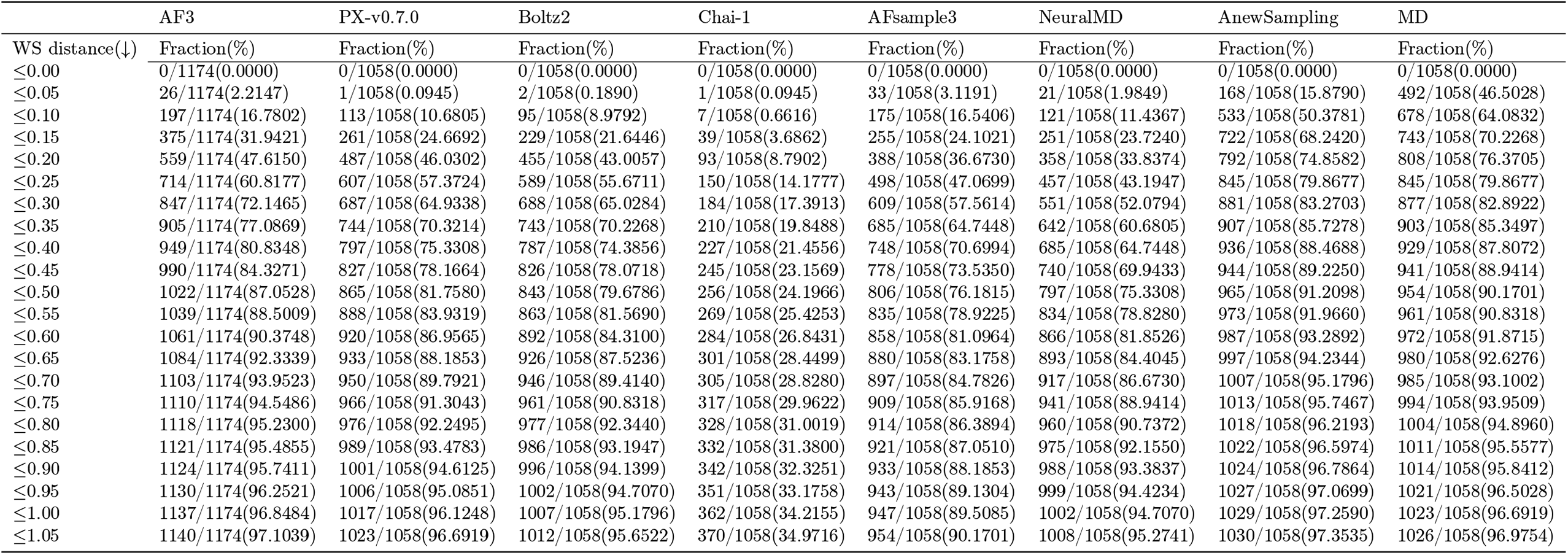
Wasserstein distance statistics for highly conserved protein-ligand interactions on the jacsmerck dataset. This table isolates structural interactions that occur with high frequency (≥ 0.8) in the reference REMD trajectories, highlighting the capacity of each model to capture strongly bound, dominant interaction states.

**Table S8.**
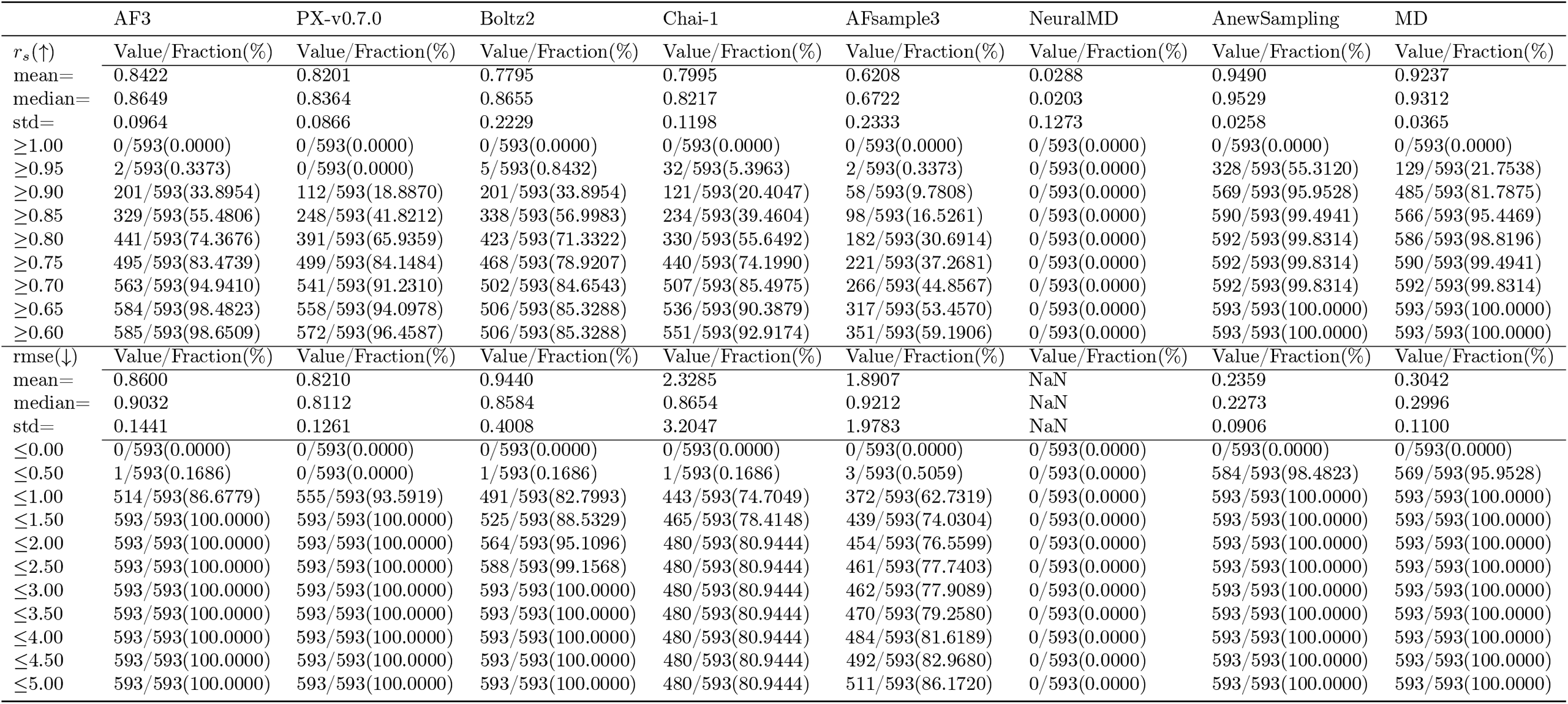
Evaluation metrics for protein backbone structural dynamics on the jacsmerck dataset. Comparative performance of each model in predicting global structural flexibility, quantified via *r*_*s*_ and RMSE of the protein backbone C*α* RMSF relative to the REMD ground truth.

**Figure S2.**
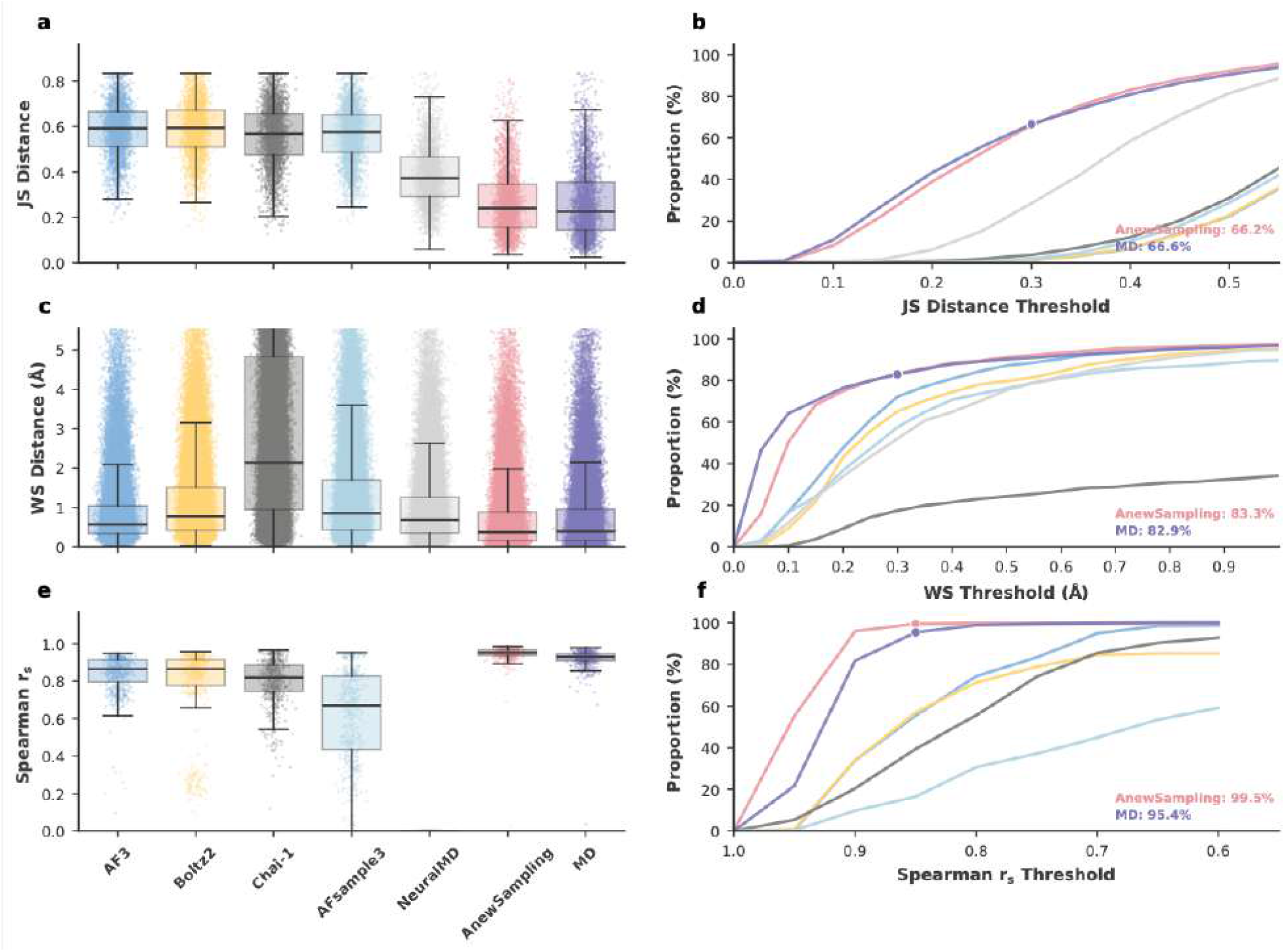
Comparative performance of generative models in capturing protein-ligand conformational dynamics for the jacsmerck dataset. **(a)** Boxplot detailing the JS distance of ligand torsion angle distributions across generated ensembles compared to the MD reference. Evaluated models include AF3, Boltz2, Chai-1, AFsample3, NeuralMD, and AnewSampling. **(b)** Cumulative proportion curves for ligand torsion JS distances. AnewSampling successfully captures ligand flexibility, achieving a 66.2% success rate that closely mirrors the MD baseline (66.6%) at the focal threshold. **(c)** Boxplot of WS distances (in Å) assessing the distributional accuracy and stability of protein-ligand interactions. **(d)** Cumulative success rate curves for interaction WS distances. AnewSampling demonstrates robust recovery of physical interaction networks, matching the rigorous MD reference (83.3% vs. 82.9%) at a stringent ≤ 0.3 Å threshold. **(e)** Boxplot evaluating the consistency of global protein backbone structural dynamics across the tested models. **(f)** Cumulative proportion curves for the *r*_*s*_ of protein backbone RMSF. AnewSampling exhibits superior preservation of coherent global motions, achieving a near-perfect 99.5% success rate compared to 95.4% for the MD baseline.

